# Meta-analysis of the human gut microbiome uncovers shared and distinct microbial signatures between diseases

**DOI:** 10.1101/2024.02.27.582333

**Authors:** Dong-Min Jin, James T. Morton, Richard Bonneau

## Abstract

Microbiome studies have revealed gut microbiota’s potential impact on complex diseases. However, many studies often focus on one disease per cohort. We developed a meta-analysis workflow for gut microbiome profiles and analyzed shotgun metagenomic data covering 11 diseases. Using interpretable machine learning and differential abundance analysis, our findings reinforce the generalization of binary classifiers for Crohn’s disease (CD) and colorectal cancer (CRC) to hold-out cohorts and highlight the key microbes driving these classifications. We identified high microbial similarity in disease pairs like CD vs ulcerative colitis (UC), CD vs CRC, Parkinson’s disease vs type 2 diabetes (T2D), and schizophrenia vs T2D. We also found strong inverse correlations in Alzheimer’s disease vs CD and UC. These findings detected by our pipeline provide valuable insights into these diseases.

**IMPORTANCE:** Assessing disease similarity is an essential initial step preceding disease-based approach for drug repositioning. Our study provides a modest first step in underscoring the potential of integrating microbiome insights into the disease similarity assessment. Recent microbiome research has predominantly focused on analyzing individual disease to understand its unique characteristics, which by design excludes comorbidities individuals. We analyzed shotgun metagenomic data from existing studies and identified previously unknown similarities between diseases. Our research represents a pioneering effort that utilize both interpretable machine learning and differential abundance analysis to assess microbial similarity between diseases.

## INTRODUCTION

The progression of many complex human diseases has been validated to be influenced by the depletion of commensal microbes associated with health, as well as the presence of potentially pathogenic microbes. The intricate interplay between the commensal microbiota and immune system has been shown to contribute to the pathogenic processes underlying various diseases (1, 2). In the context of standard microbiome studies, most diseases are typically studied in isolation. They are compared against a control group that is constructed to minimize confounding factors (3). This approach, however, neglects the fact that many patients have multiple comorbidities (4). Currently, nearly 60% of adult Americans have at least one chronic disease, with about 40% having multiple conditions (5). In medicine, it is common to study the nature of the comorbidities to understand the etiology of a given disorder. Within the field of oncology, genetic sequencing has been used to unravel similarities between disorders that were not observed from traditional medical observation (6). Doing so has provided a scaffold for drug-repositioning to repurpose drugs to target similar cancer types.

We propose that the same approach should be taken when conducting microbiome studies. Existing literature also offers strong motivation for undertaking this study. Firstly, microbes function as a central hub for the metabolism of dietary compounds, thereby serving as a coordination point for the distribution of nutrients (7). Consequently, it is plausible that any metabolic disorder may have an associated microbiome component (8). Secondly, microbes have a strong interaction with the immune system (9), thus play a critical role in the development and modulation of the immune system. Thirdly, microbes are known to metabolize ingested drugs, and are known to contribute to efficacy (10, 11). All these facts indicate that microbes can have a multifaceted impact on disorders that have not been previously linked to the gut microbiome. Consequently, it is highly possible that there are disorders that are observed to have similar microbiome patterns, despite having dissimilar disease phenotypes.

Previous studies have shown that the imbalance of the bacterial community played a contributory role in the development of complex disorders, including neurological disorders, immune disorders, metabolic disorders, and gastrointestinal disorders. Recent studies have further highlighted the shared microbial signatures that contribute to these diseases, underscoring the need for more in-depth studies. For instance, *Prevotella copri* was found to be more prevalent in both type 2 diabetes (T2D) and rheumatoid arthritis patients compared to healthy controls, possibly due to its immune-relevant role in the pathogenesis (12, 13). More recently, dysregulation of the gut-brain axis (GBA) has been demonstrated to contribute to the development of several neurological disorders, such as Alzheimer’s disease (AD), autism spectrum disorder (ASD) and mood disorders (14, 15). We explored the intersection of microbial signatures associated with those disorders, leveraging existing datasets that delve into the gut microbiome of these conditions.

Our goal is to provide a computational pipeline that can measure disease similarity based on microbiome composition. We utilize interpretable machine learning and differential abundance methods to identify both disease-specific microbes and microbes that are commonly observed across diseases. The key to comparing multiple disease cohorts was leveraging recent insights from removing batch effects within studies (15, 16). While previous meta-analysis research has mainly focused on analyzing multiple diseases to understand what makes each disease unique (17, 18), our pipeline represents the largest shotgun metagenomic meta-analysis conducted to measure the similarity between diseases with high resolution. We focus on diseases that are found to be associated with the imbalance of gut microbiome. We included datasets investigating 11 disorders ranging from metabolic disorders, gastrointestinal (GI) disorders, neurological disorders to cancer.

Since estimating disease similarity is a necessary first step prior to drug repositioning, we provide a modest first step in highlighting the possibility of incorporating microbiome insights into the drug-repositioning pipeline. To investigate the similarity between these disorders, we focus on shotgun metagenomics. While there is a lot more 16S rRNA gene data that is available, we have opted not to include these due to the lack of species resolution and genomic insights. This allows us to obtain high level species or strain resolution while gaining insights to the potential functional roles of these microbes. We address a critical gap in understanding complex disease by examining shared microbial signatures.

## RESULTS

We have developed a novel pipeline (Figure 1) that computes disease similarity at both microbial species and gene level, enabling a consistent data standard to make different studies more comparable. We compiled a large multi-study meta-analysis, with consistent processing to enable comparisons across studies that accounts for batch effects. Our findings reveal a high degree of similarity between Crohn’s disease (CD) vs ulcerative colitis (UC), CD vs colorectal cancer (CRC), Parkinson’s disease (PD) vs T2D, as well as schizophrenia vs T2D. Our results show that the similarity at the microbial species level was consistent with the similarity at the microbial gene level, explained by both the enrichment of pathogenic microbes and the depletion of beneficial microbes. Lastly, we found that the microbial gene profiles between AD and inflammatory bowel disease (IBD) are anticorrelated, highlighting a more pronounced metabolic distinction between these two disorders than previously suspected.

**Figure 1.**
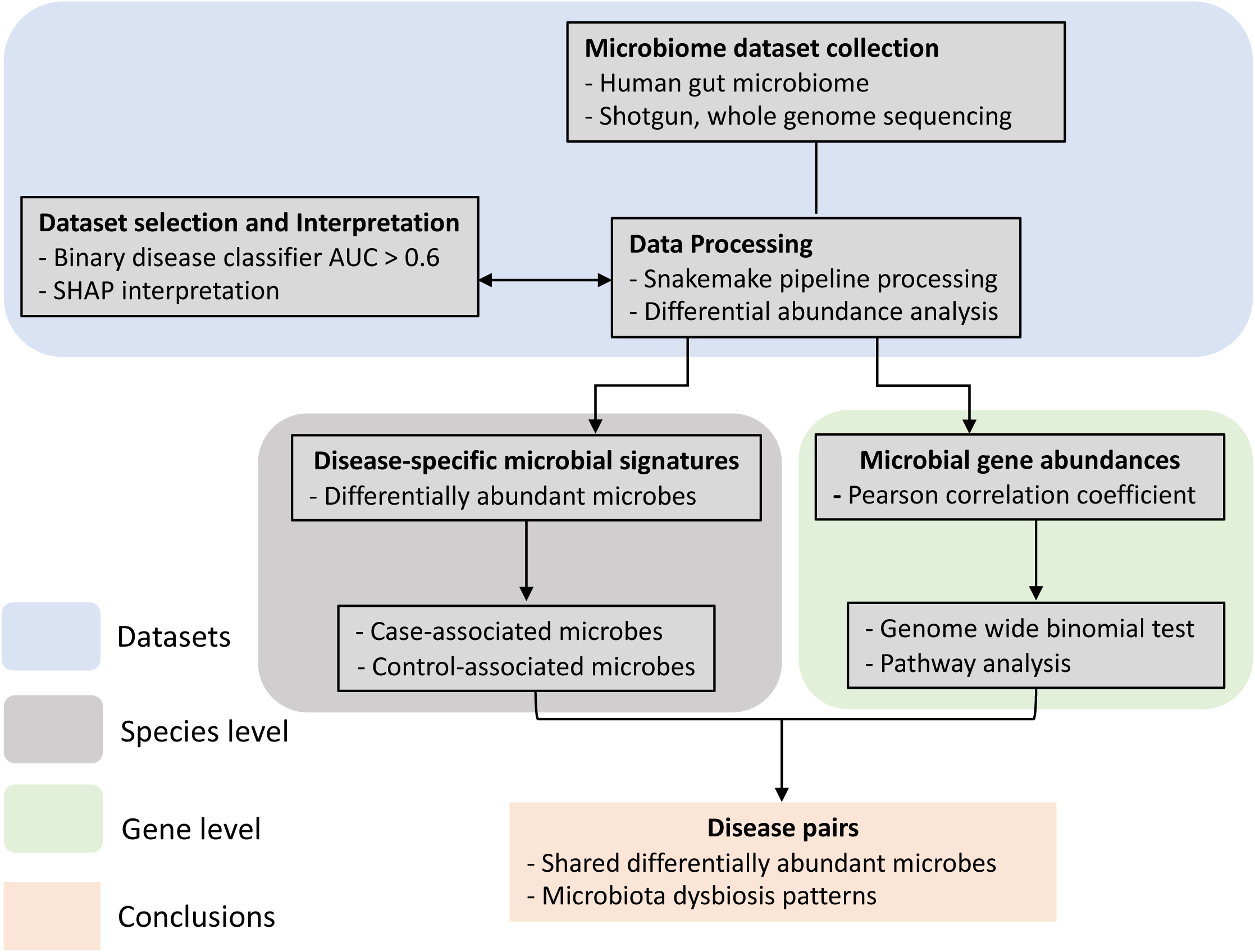
The overall design and data analysis pipeline. Flow Chart of this meta-analysis. First, shotgun metagenomic datasets investigating the human gut microbiome in multiple diseases were curated and processed consistently with the Snakemake metagenomic pipeline built in this study, and the microbial abundances matrices were generated. Second, Gradient Boosting and Random Forest classifiers were built for each dataset/disease. Then datasets with classification accuracy above the threshold of 0.6 remained in the following analysis. Disease specific microbial signatures and microbial similarity at the species level were analyzed with the differential abundance results. At the gene level, Pearson correlation coefficients of microbial genes between every disease pair were calculated and used as the proxy for disease similarity. Disease pairs that showed high or low similarity were further investigated with pathway analysis.

### Consistent data processing and cohort selection for this meta-analysis

In this study, we applied the Snakemake pipeline to process available samples, and constructed binary disease classifiers for each study. After fitting both binary gradient boosting (GB) and random forest (RF) classifiers, we found that GB classifiers showed better overall performance across diseases. We then employed GB classifiers in subsequent studies and utilized them to exclude studies that cannot discriminate the disease phenotype based on microbial profile. The resulting dataset derived from 18 studies (19–35), encompasses a total of 2091 samples (Table 1). These samples are distributed across 11 countries spanning Europe, Asia, and America. The following diseases were included: four neurological disorders (AD, ASD, schizophrenia, and PD); two autoimmune disorders (multiple sclerosis (MS) and type 1 diabetes (T1D); two metabolic disorders (obesity and T2D); two GI disorders (CD and UC); and one cancerous disorder (CRC). We compared the classifier performances per cohort and per disease. Per cohort refers to build classifiers for each dataset, and per disease refers to build classifiers with the datasets for a disease combined. The results showed that the overall classification accuracy increased when it was tested per disease. Suggesting that the consolidation of datasets from diverse cohorts enhances the overall representativeness of the disease, consistent with previous findings investigating both ASD (15) and CRC (26).

**Table 1.**
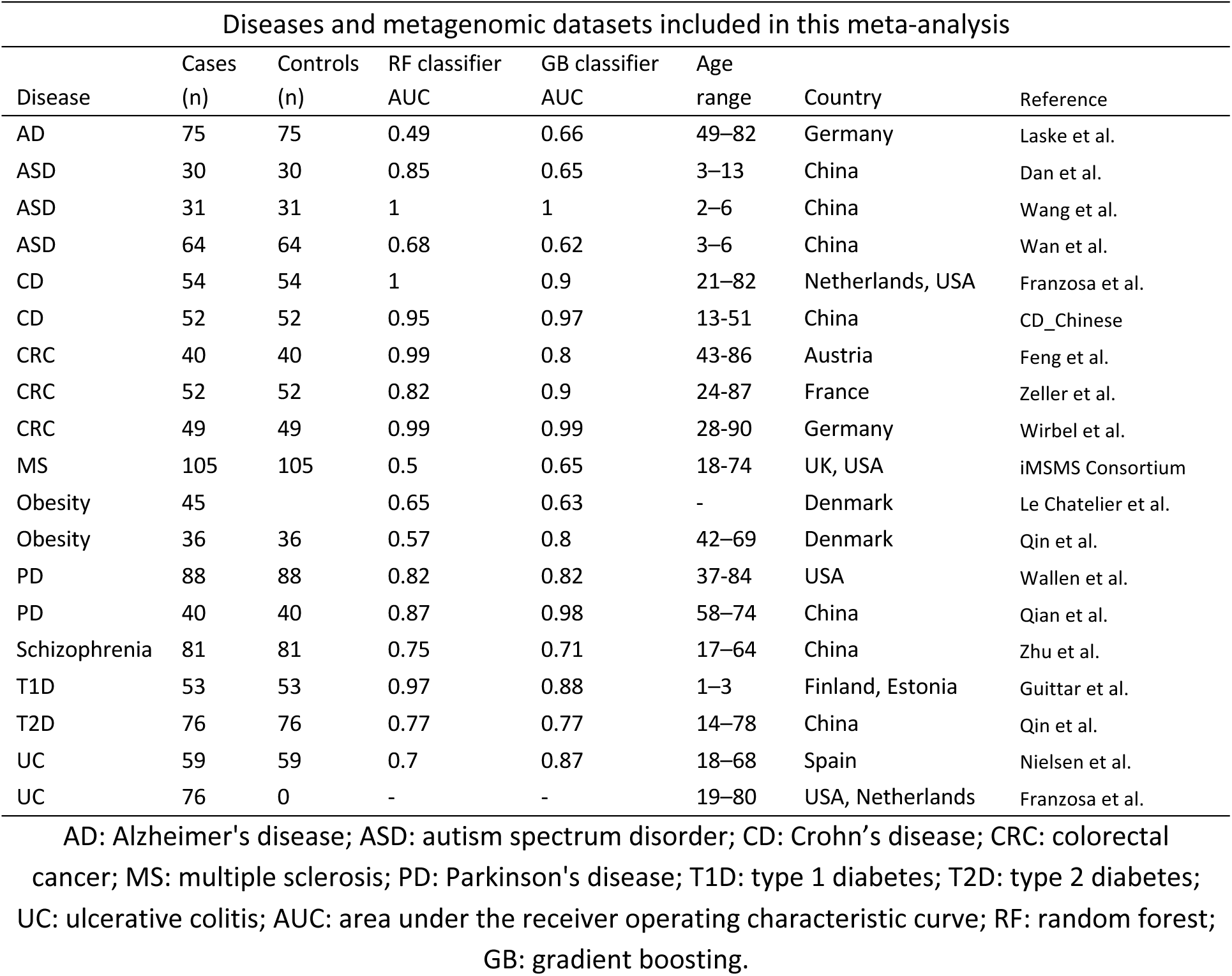
Diseases and metagenomic datasets included in this meta-analysis.

| Diseases and metagenomic datasets included in this meta-analysis |  |  |  |  |  |  |  |
| --- | --- | --- | --- | --- | --- | --- | --- |
| Disease | Cases<br>(n) | Controls<br>(n) | RF classifier<br>AUC | GB classifier<br>AUC | Age<br>range | Country | Reference |
| AD | 75 | 75 | 0.49 | 0.66 | 49–82 | Germany | Laske et al. |
| ASD | 30 | 30 | 0.85 | 0.65 | 3–13 | China | Dan et al. |
| ASD | 31 | 31 | 1 | 1 | 2–6 | China | Wang et al. |
| ASD | 64 | 64 | 0.68 | 0.62 | 3–6 | China | Wan et al. |
| CD | 54 | 54 | 1 | 0.9 | 21–82 | Netherlands, USA | Franzosa et al. |
| CD | 52 | 52 | 0.95 | 0.97 | 13–51 | China | CD_Chinese |
| CRC | 40 | 40 | 0.99 | 0.8 | 43–86 | Austria | Feng et al. |
| CRC | 52 | 52 | 0.82 | 0.9 | 24–87 | France | Zeller et al. |
| CRC | 49 | 49 | 0.99 | 0.99 | 28–90 | Germany | Wirbel et al. |
| MS | 105 | 105 | 0.5 | 0.65 | 18–74 | UK, USA | iMSMS Consortium |
| Obesity | 45 |  | 0.65 | 0.63 | - | Denmark | Le Chatelier et al. |
| Obesity | 36 | 36 | 0.57 | 0.8 | 42–69 | Denmark | Qin et al. |
| PD | 88 | 88 | 0.82 | 0.82 | 37–84 | USA | Wallen et al. |
| PD | 40 | 40 | 0.87 | 0.98 | 58–74 | China | Qian et al. |
| Schizophrenia | 81 | 81 | 0.75 | 0.71 | 17–64 | China | Zhu et al. |
| T1D | 53 | 53 | 0.97 | 0.88 | 1–3 | Finland, Estonia | Guittar et al. |
| T2D | 76 | 76 | 0.77 | 0.77 | 14–78 | China | Qin et al. |
| UC | 59 | 59 | 0.7 | 0.87 | 18–68 | Spain | Nielsen et al. |
| UC | 76 | 0 | - | - | 19–80 | USA, Netherlands | Franzosa et al. |

AD: Alzheimer's disease; ASD: autism spectrum disorder; CD: Crohn's disease; CRC: colorectal cancer; MS: multiple sclerosis; PD: Parkinson's disease; T1D: type 1 diabetes; T2D: type 2 diabetes; UC: ulcerative colitis; AUC: area under the receiver operating characteristic curve; RF: random forest; GB: gradient boosting.

Our analysis revealed three diseases with with-in cohort cross-validation Area Under the receiver operating characteristic Curve (AUC) exceeding 0.95 utilizing different machine learning algorithms in certain datasets: ASD, CD, and CRC (Table 1). Notably, both CD and CRC are diseases related to GI, predominantly impacting the GI tract. Within the ASD cohort, where we observed high classification accuracy, most of the ASD patients also have GI symptoms (36). Classifier is trained on the training dataset, and its predictive accuracy is assessed on a hold-out test dataset. This is important to emulate real-world clinical environments, where there could be drift between clinical studies due to confounding demographics and experimental protocols. Additionally, we collected a hold-out cohort evaluation using two independent datasets to assess the generalization performance of the binary classifiers on previously unseen test datasets for CD (35) and CRC (37).

### Comparison of Crohn’s disease and colorectal cancer: SHAP (SHapley Additive exPlanations) interpretation and differential abundance analysis

Population-based cohort studies have found that CD is a risk factor for CRC (38), we choose to compare the shared microbial signature between CD and CRC first as a sanity check for our analysis. We applied two distinct approaches to gain insights. Firstly, we used SHAP to interpret the binary disease classifiers. SHAP provides a valuable means of understanding the contribution of each feature in the classification process, offering interpretability to complex machine learning models. Secondly, we conducted a comprehensive analysis of differential abundance. This approach allows us to identify significant variations in the abundance of microbes between disease cases and healthy controls. Since these two quantities are generated from distinct methods, they provide different perspectives that are sometimes in conflict. By leveraging both pieces of information, we looked at microbes that could be strongly explained by both the Shapley values and large log-fold changes. While we did not observe an overlap in the features that contribute most to the classification for CD and CRC, we found there is a considerable overlap in the microbial species that exhibit differential abundance in CD and CRC patients.

Both the binary classifiers of CD and CRC displayed robust generalization abilities when tested on previously unseen cohorts, with the AUC values of 1.00 and 0.87, respectively. In the case of CD classification, control-associated species within the *Faecalibacterium* genus, such as *F. sp3900551435* (shapley rank 1st in CD control and rank 2nd in CD case) and *F. prausnitzii* (shapley rank 6th in CD control and rank 7th in CD case), exhibit high absolute shapley values in both cases and controls of the CD cohort (Supplementary Table 1). As demonstrated in prior research, *F. prausnitzii* can produce proteins with anti-inflammatory properties and is involved in CD pathogenesis (39). Altogether, our results implies that these control-associated microbes played a substantial role in distinguishing CD patients from controls (Figure 2a).

**Figure 2.**
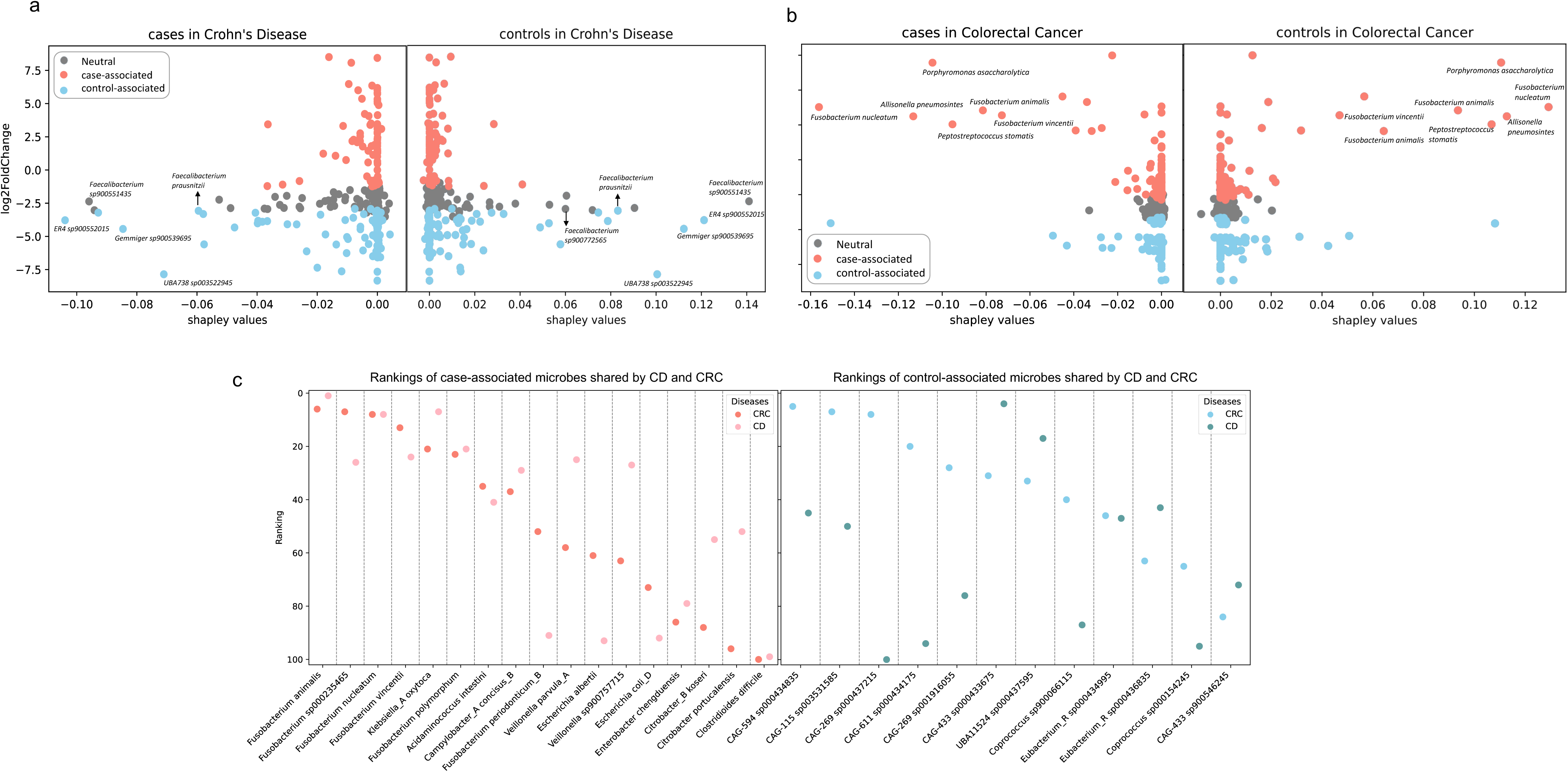
Interpretation of binary classifiers and differentially abundant microbes overlaps in Crohn’s Disease (CD) and Colorectal Cancer (CRC). The differential abundant microbes are identified first by computing LFC between case and control within one disease, then ranked by the 5% Confidence Interval (CI) of LFC to identify the top-100 case-associated microbes, finally ranked by the 95% CI of LFC to identify the top-100 control-associated microbes. Each dot represents one microbe, and its color is coded by its ranking. Dots colored blue and salmon represent the microbes differentially abundant in disease cases and controls respectively. Dots colored gray are the ones that are considered neutral. (a)-(b) X axis is the shapley values, Y axis is the log2 fold change (LFC) between case and control. Left panels are the cases, which have a sum of shapley values as negative values, right panels are the controls, which have a sum of shapley values as positive values. Dots with high absolute Shapley values and high log2FC are labeled. (a) Shapley values vs LFC in CD cases and controls. (b) Shapley values vs LFC in CRC cases and controls. (c) Overlap of the differentially abundant microbes between CD and CRC. X axis is the microbes, y axis is the microbe’s rankings. A smaller ranking number for case-associated microbes indicates a greater increase of the microbe in disease cases. A smaller ranking number for control-associated microbes indicates a greater increase of the microbe in healthy controls.

However, in CRC, case-associated microbes had a more pronounced influence on CRC classification (Figure 2b), particularly exemplified by *Fusobacterium nucleatum*, *Allisonella pneumosintes*, and *Prophyromonas asaccharolytica* (Supplementary Table 2). Specifically, *F. nucleatum* was ranked first in terms of shapley values in both CRC case and control groups. *F. nucleatum* was known to be enriched in colorectal adenomas and adenocarcinomas (40) and it can create a proinflammatory microenvironment that supports the progression of colorectal neoplasia (41). It’s also one of the common oral bacteria. This has been previously observed in lung cancer, where oral commensals are more abundant in the lower airway of lung-cancer patients compared to the control population (42). Recent studies found the connection between oral bacteria and gut is possibly through both the ectopic gut colonization by oral commensals and induction of migratory Th17 cells, constituting a complex interplay between the microbiome and immune system (43, 44). Our results confirmed its pivotal role in distinguishing CRC patients from controls. We also identified other candidates that warrant further investigation.

It has been found that individuals diagnosed with CD may face an elevated risk of developing CRC, possibly due to the chronic inflammation associated with CD (38, 45). Our findings revealed the overlapped case-associated microbes between CD and CRC contributed to the similarities of these two diseases, such as *Fusobacterium spp.* and *veillonella spp*. along with the shared depletion of potential probiotic *Coprococcus spp.* (Figure 2c). It is worth noting that we found many *Fusobacterium* species worth looking into (*F. animalis, F. sp000235465, F. nucleatum, F. vincentii, F. polymorphum*), including one of them (*F. prausnitzii*) that have been validated by previous studies (39). Differential abundance analysis found 19% and 17% overlap of the case-associated and control-associated microbes respectively between these two diseases (Figure 3a). Our findings, in agreement with previous studies, highlight the potential involvement of these shared microbial signatures in the progression of both CD and CRC. The shared microbial features we find here are crucial for us to better understand the common features between these diseases, and can help us step closer to the real therapeutic target for these complex diseases.

**Figure 3.**
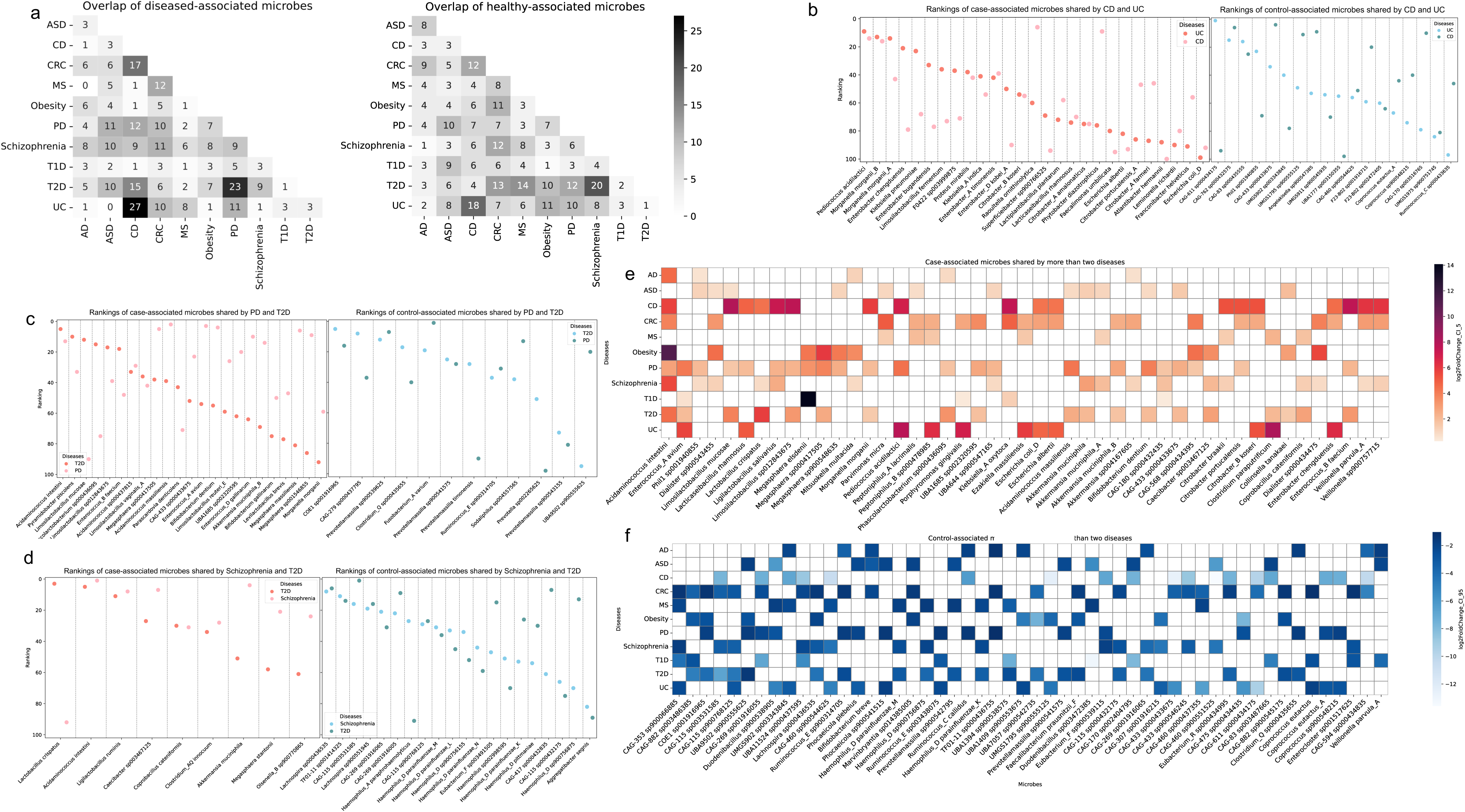
Microbial species level similarity between diseases. (a) The annotation numbers represent the number of microbes overlapping between two diseases among the top 100 case-associated microbes or the top 100 control-associated microbes. (b)-(d) Dots colored in salmon represent case-associated microbes and their rankings. A smaller ranking number indicates a greater increase of the microbe in disease cases. Dots colored in blue represent control-associated microbes and their rankings in controls. A smaller ranking number indicates a greater increase of the microbe in healthy controls. (b) Overlap of the differentially abundant microbes between Crohn’s Disease (CD) and Ulcerative Colitis (UC). (c) Overlap of the differentially abundant microbes between Schizophrenia and Type 2 Diabetes (T2D). (d) Overlap of the differentially abundant microbes between Parkinson’s Disease (PD) and T2D. (e) (f) Differential abundant microbes shared by more than two diseases. (e) Case-associated microbes shared by more than two diseases. (f) Control-associated microbes shared by more than two diseases.

### Differential abundance analysis revealed disease pairs with high similarity at the microbial species level

Among the top disease-pairs exhibiting the most significant overlap of case-associated microbes, CD vs UC had the highest co-occurrence, followed by PD vs T2D (Figure 3a). For case-associated microbes, 27% were common to both CD and UC, 23% were shared between PD and T2D. On the other hand, schizophrenia vs T2D, as well as CD vs UC, showed a substantial overlap of control-associated microbes. Specifically, 20% and 18% of the top control-associated microbes were shared in these pairs, respectively (Figure 3a). We choose to conduct a more in-depth comparison at microbial species level for those disease pairs: CD vs UC, PD vs T2D, and schizophrenia vs T2D. Based on the observation that these pairs exhibited the highest overlaps of differential abundant microbes. We identified the shared differential abundant microbes for these three disease pairs. Differential abundant microbes shared by more than two diseases have also been identified in our study (Figure 3).

Among the case-associated microbes shared by CD and UC, recognizable microbes such as *Pediococcus acidilactici*, and known commensal species like *Morganella morganii* (Figure 3b, 3e), are potentially involved in gut dysbiosis. A recent study in mice has confirmed that the overabundance of *P. acidilacitic* may play a role in triggering IBD by producing lipopolysaccharide and exopolysaccharide byproducts (46). Cao *et al.* have demonstrated that *M. morganii* isolated from IBD patients can generate genotoxic metabolites called indolimines. These metabolites have the potential to induce DNA damage and contribute to cancer progression (47). Within the control-associated microbes, *Coprococcus eutactus*, a potent probiotic that can alleviate colitis through acetate-mediated IgA response and microbiota restoration (48), is amongst the most prevalent in the control populations. Our results unveiled several other case-associated microbes that warrant further investigation, including species from the genera *Enterobacter* and *Citrobater*, among others (Figure 3b). Altogether, this highlights the strong microbial similarity between the two IBD subtypes, which is consistent with both previous microbiome studies and the clinical phenotype (23, 49).

We found control-associated microbes depleted in both schizophrenia and T2D include species from *Lachnospira* and *Haemophilus* genera (Figure 3c, 3f). Microbes from the Lachnospiraceae family are known to produce butyrate, which is one of several SCFAs that has beneficial effects on cellular metabolism and intestinal homeostasis. The loss of such microbes is linked to chronic inflammation and is likely involved in metabolic diseases such as T2D (50). On the other hand, *Haemophilus parainfluenzae* is a common commensal that has been recognized as an opportunistic pathogen, but its specific functional role remains unclear. Studies have reported a lower abundance of *H. parainfluenzae* in mental disorders compared to healthy controls (51). We found the decreased abundances of these microbes contribute to the similarity between schizophrenia and T2D.

Similar to schizophrenia, PD is another neurological disorder that showed high similarity with T2D, however mainly contributed by the shared increase of case-associated microbes in patients (Figure 3d). We observed that *M. morganii* is also differentially increased in both PD and T2D patients, along with species from the genera *Acidaminococcus*, *Limosilactobacillus*, and others. The increased abundance of *Acidaminococcus intestini* in disease cases has been found in seven diseases (Figure 3e), making it the most commonly shared case-associated microbe in our analysis. A recent cross-sectional study found *Acidaminococcus intestini* was one of the microbes that were more abundant in subjects consuming the most pro-inflammatory diets (52). We did observe that *Akkermansia muciniphila* was differentially increased in all these disorders (PD, schizophrenia, and T2D) as well (Figure 3e). This finding is highly controversial in the literature, since previous studies have observed that *Akkermansia muciniphila* is both beneficial (53) and pathogenic (54). From our analysis, it is difficult to determine the causal role of *Akkermansia muciniphila* in these diseases. Follow up mechanistic and clinical studies will be necessary to explore the involvement of this microbe in more depth.

### Microbial gene level comparison and pathway analysis showed consistency with species level results

Disease similarity based on the microbial genes can be accessed by comparing the Pearson correlation coefficient R between the inferred log2 fold changes (LFC) across every two diseases (Figure 4a). The R value for CD vs UC stands at 0.6, representing the highest positive correlation observed across all disease pairs (Figure 4b). As two subtypes of IBD, both CD and UC are characterized by transmural inflammation, with CD being able to affect any area from the mouth to the perianal region while UC is limited to the colon’s mucosal layer (55). Previous studies have demonstrated that IBD is influenced by genetic predisposition, immune system dysregulation, and environmental factors (23, 35). Our pathway analysis revealed the involvement of both case-associated and control-associated microbes in various metabolic pathways. Specifically, we identified discrepancies in amino acid metabolism, energy metabolism, and lipid metabolism that differentiated between case-associated microbes and control-associated microbes for CD and UC. Case-associated microbial genes exhibited a higher prevalence within most of these metabolic pathways, many of which contributed to inflammation and infection (Figure 4e). Pathogenic microbes such as *Fusobacterium*, *Klebsiella,* and *Stenotrophomonas* (56–58) are heavily involved in pathways including tryptophan biosynthesis, oxidative phosphorylation, and fatty acid biosynthesis. This is consistent with previous studies, highlighting the microbial and clinical similarity between these two disorders.

**Figure 4.**
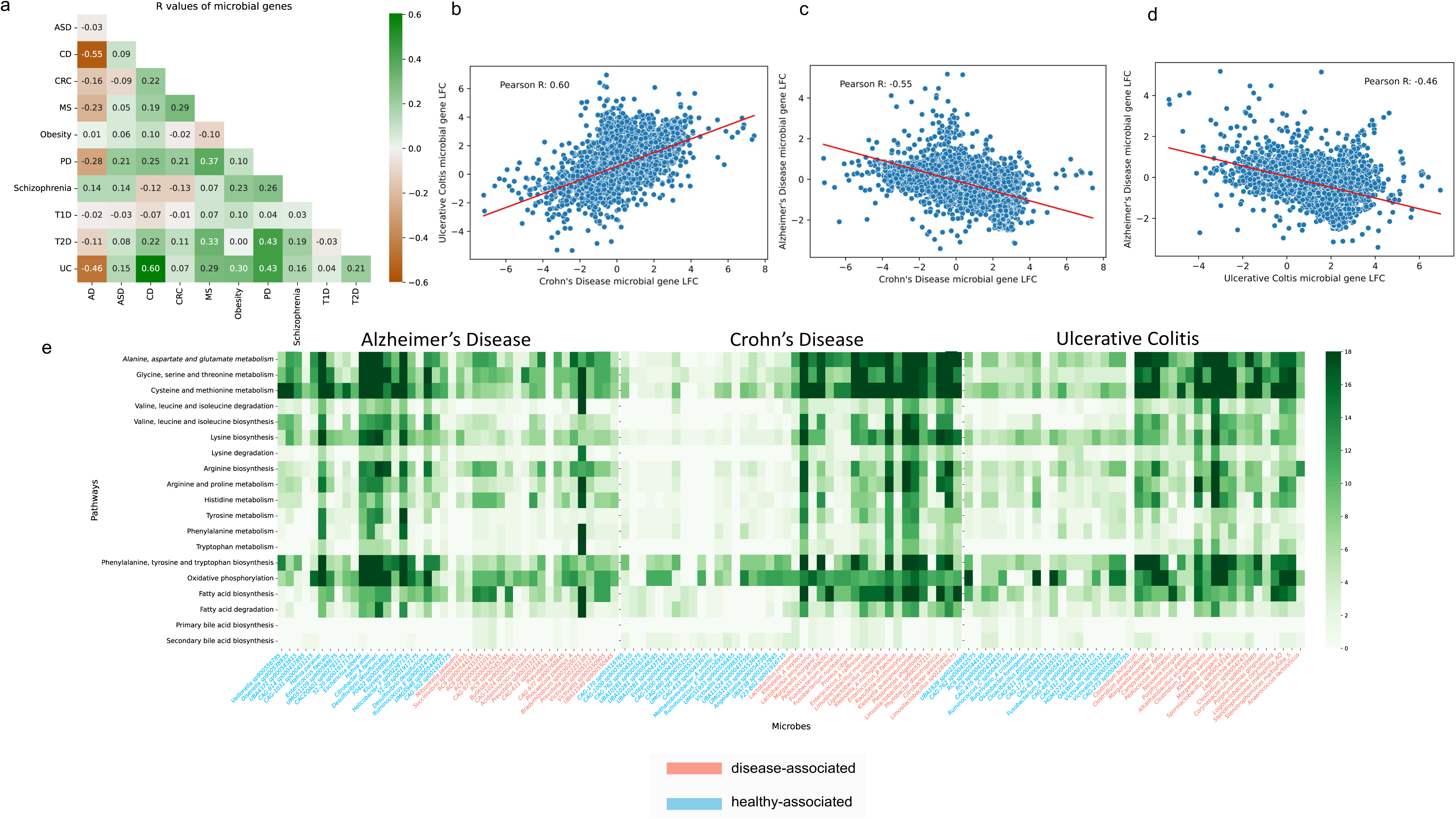
Microbial gene level similarity between diseases and the pathway signatures of the microbes. (a) Pearson correlation coefficient R between the inferred microbial gene log2 fold changes across every two diseases. (b) Scatterplot of the Pearson R between Crohn’s Disease (CD) and Ulcerative Colitis (UC). (c) Scatterplot of the Pearson R between CD and Alzheimer’s Disease (AD). (d) Scatterplot of the Pearson R between UC and AD. (e) Amino acid metabolism, energy metabolism, and lipid metabolism pathways of the microbial signatures in AD, CD, and UC. The x axis are the differential abundant microbes, the blue ones represent control-associated microbes, while the salmon ones represent the case-associated. The y axis are the KEGG pathway modules. The numbers on the right green bar represent the number of genes.

Conversely, AD has a strong negative correlation between differential gene abundances between both CD (R=-0.55) (Figure 4c) and UC (R=-0.46) (Figure 4d), highlighting how the microbial link with AD affects the same pathways, but possibly through a different mechanism of action. Amongst the genes that were most differentiating between cases and controls for CD, UC, and AD, many of these genes are involved in the pathways related to the metabolism of amino acid, lipid, and energy metabolic. In AD patients, this was partially due to the decrease in microbes, such as *Hafnia alvei*, *Ruminococcus*, and *Citrobacter*, which had a greater prevalence of genes that are encoded in these pathways (Figure 4e). Some of the control-associated microbes are known to generate metabolites like histamine, conjugated fatty acids, and dopamine, which act as neuroprotective agents in AD (59). AD is known for the accumulation of beta-amyloid plaques and tau tangles in the brain (19), and is often characterized by metabolic abnormalities, including compromised bioenergetics, impaired lipid metabolism, and an overall decreased metabolic capacity (60).

Most of the drugs designed to treat AD patients, such as Lecanemab, Donanemab, and Remternetug, are focused on removing plaque (61). In contrast, many drugs that target IBD are immunosuppressants, such as Azathioprine, Mercaptopurine (6-MP), and Methotrexate (62). While drugs used to treat IBD and AD are known to have very different functional roles, it is interesting to see how the microbial gene profiles between these two disease populations have discordance in the same metabolic pathways (Figure 4e). It is currently not clear to us why this discordance exists. However, these findings highlight interesting directions for pre-clinical follow up studies, particularly in exploring the utility of immune-enhancing drugs in AD.

### Both the enrichment of pathogens and depletion of control-associated microbes contribute to the similarity between complex human diseases

Various types of microbiome shifts in complex human diseases have been identified by previous studies, encompassing the depletion of beneficial microbes, enrichment of pathogens, and a comprehensive reconstruction of gut microbial communities (17). In many of the disease pairs that exhibit a high overlap in microbial signatures, we found that both the enrichment of pathogens and the depletion of beneficial microbes contribute substantially to their similarity. This holds true regardless of the previous classification of their dysbiosis patterns in prior studies.

Dysbiosis associated with CRC was generally characterized by increased prevalence of the pathogenic microbes (25) while CD was consistently characterized by the depletion of control-associated microbes (63). Combined similarity networks with the sum of overlapped microbe weights show that both shifts contribute to the similarities between diseases (Figure 5). The color of edges shows the difference of shifts, and the width of edges between two diseases are proportional to the overlapped microbes. The similarities between CD and CRC comprise a mixture of both shifts. This indicates that the dysbiosis patterns of some diseases are more complicated than initially clarified, opening new opportunities for repurposing narrow-spectrum antimicrobials and probiotic treatments.

**Figure 5.**
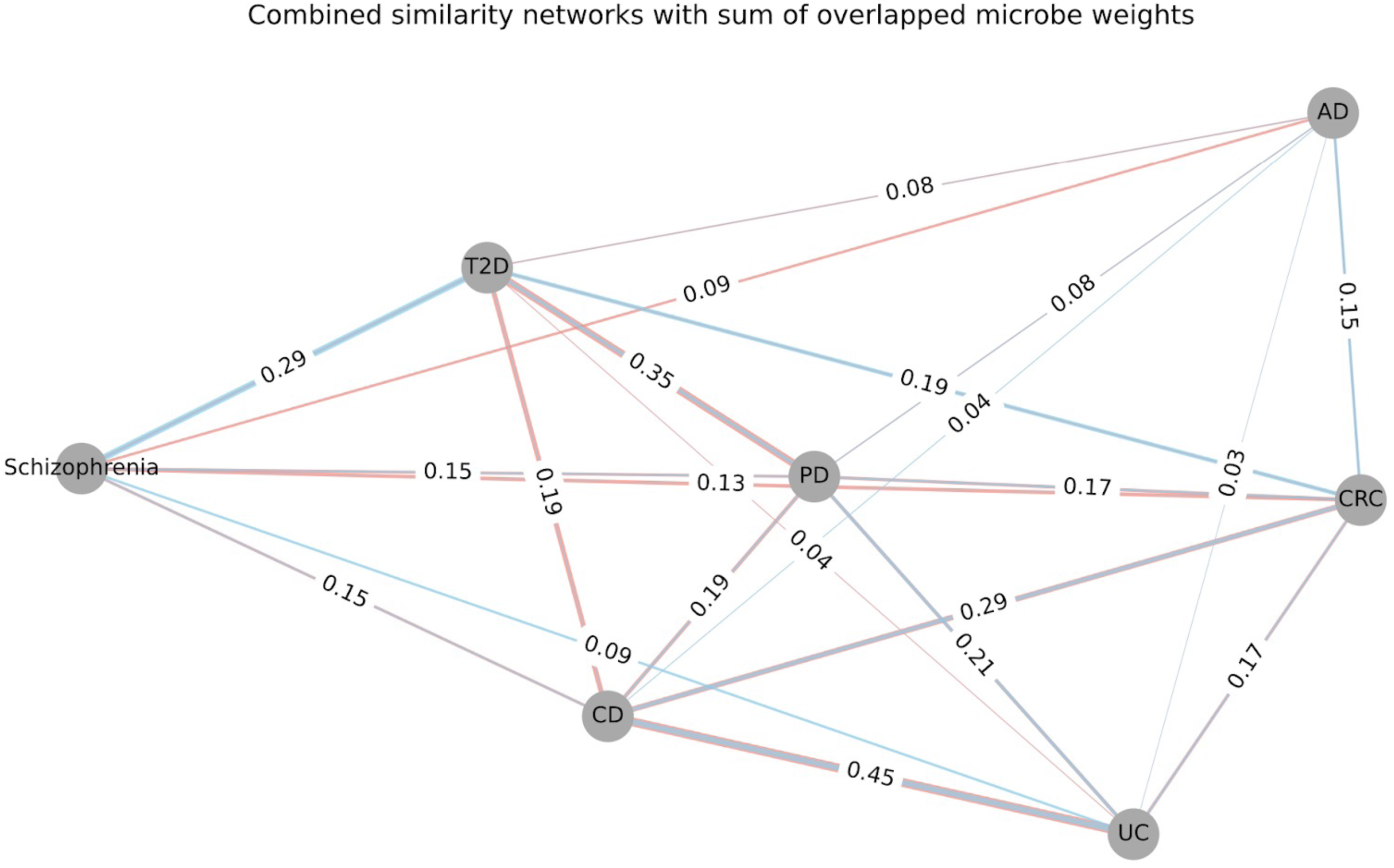
Combined similarity networks with sum of overlapped microbe weights. Each node represents one disease type, the weight of edges shows how similar two diseases are. The number in each edge is proportional to the overlapped differential abundant microbes in each disease (case vs control): top-100 (case-associated) and bottom-100 (control-associated). The colors of the edges indicate the origin of the similarities: salmon color edges represent the similarity conferred by the overlap of case-associated microbes; the blue color represents the similarity conferred by the overlap of control-associated microbes.

There is consistency between the similarity observed at the microbial species level and that at the microbial gene level. AD and the two IBD subtypes showed the least overlap in differentially abundant microbes. They also exhibited the least similarities at the microbial gene level. Furthermore, disease pairs like CD vs UC, CD vs CRC, PD vs T2D, as well as schizophrenia vs T2D, demonstrated a high overlap in differentially abundant microbes and high Rs in microbial gene abundances. Discrepancies may arise when comparing these similarities at different levels. For instance, both PD vs UC and PD vs T2D have strong microbial gene similarities (R=0.43, and R=0.43) (Figure 3a). However, PD vs UC has a very small overlap in differentially abundant microbes (overlap=21%), while PD vs T2D has a larger overlap (overlap=35%) (Figure 5). This is consistent with the functional redundancies that have been observed in microbial communities, even if there is small overlap between the microbial taxa, there could still be strong overlap in the metabolic function due to the common metabolic roles different microbes play (64).

## DISCUSSION

We have assembled the largest shotgun metagenomics meta-analysis that has inferred disease similarity with high resolution, across 2091 samples from 18 studies, encompassing 11 different disease types. We conducted case-control differential abundance analysis within each disease, following comparison between diseases. Our results demonstrated that binary disease classifiers for CD and CRC exhibit a strong generalization capability when applied to unseen data. We discovered a high degree of microbial similarity between CD and CRC. This finding aligns with the fact that CD is a risk factor for CRC, thereby validating our pipeline. Furthermore, CD and UC are detected to have the strongest microbial similarity. Given that both are subtypes of IBD, this observation further substantiates the effectiveness of our pipeline.

We identified two neurological disorders, PD and schizophrenia, that exhibited high microbial similarity with T2D. Higher prevalence of T2D in schizophrenia patients has been observed in observational clinical studies (65), but there has not been a microbiome connection that has been previously established. Our findings offer valuable perspective on the potential for repositioning T2D drugs to treat these neurological disorders or vice versa. Metformin is a commonly used oral treatment for T2D (66). Metformin alters the gut microbiome of T2D patients and altered gut microbiota mediates some of metformin’s antidiabetic effects (67). Interestingly, recent studies suggest that metformin has a positive effect on conditions such as anxiety or depression (68, 69). Mouse study also established the neuroprotective effect of metformin in PD and supports the therapeutic potential of metformin in the treatment of PD (70, 71).

One surprising finding we discovered was that microbiome profiles in AD were anti-correlated with microbiome profiles in IBD. Microbiome components have been observed to play a role in both diseases. In a study on AD involving fecal microbiota transplantation (FMT), fecal microbiota from Alzheimer’s patients and age-matched healthy controls were transplanted into microbiota-depleted rats respectively. It was observed that the severity of impairments in hippocampal neurogenesis in these rats correlated with the clinical cognitive scores of the donor patients (72). Multiple FMT in IBD have shown promising results reducing inflammation in patients (73, 74). However, common biological mechanisms between these two diseases have not been previously established. AD is characterized by inflammation in the brain, while IBD is mainly characterized by GI inflammation. The anti-correlated microbial gene profiles between AD and IBD highlights potentially novel directions for drug design. If drugs designed to target IBD were applied to Alzheimer’s patients, would it antagonize Alzheimer’s symptoms? Furthermore, is it possible for these efforts to uncover new therapeutic strategies that could counteract the effects of these drugs?

While our findings provide valuable insights, it is important to note that our study is subject to several notable limitations. First, there are multiple confounding factors that could bias our findings. For instance, most studies did not perform absolute quantification, thus it is not possible to identify microbes that are truly differential between the case and control populations (75). It is possible that the microbes detected was due to our choice of reference frame, we assume that the average microbe isn’t changed between the case-control cohorts, but if there is a significantly altered microbial load between the cohorts, that could lead to false positives or false negatives in the differential abundance results (75). To ameliorate this issue, we focused on the top 100 microbes differentially increased in the cases and the top 100 microbes differentially increased in the controls to avoid the issue of identifying an unstable reference frame. There are also likely a few biological confounders that aren’t well-documented but could affect our findings, such as medication history (i.e., antibiotics usage) or dietary patterns.

To improve our ability to perform causal inference, it is important to not only account for these relevant confounders, but also take advantage of longitudinal observational cohorts and clinical trials to identify indirect effects on outcomes due to external interventions. Incorporating multiple omics levels will also help improve causal resolution, since increasing the number of observed biomarkers will increase the chances of observing a biomarker that plays a causal role in the disease symptoms. Our analysis focused solely on shotgun metagenomics data, overlooking the potential insights offered by other omics-level data (76). Host transcriptomic profiles facilitated the identification of host gene-microbiome associations in gastrointestinal disorders (77). Metabolomics would yield insights into lipid and bile-acid metabolism which has been observed in the context of IBD (78). Proteomics will likely play an important role in understanding amino acid metabolism and immune response, which we have shown to play a role in estimating disease similarity. While observational and clinical studies can help identify putative causal biomarkers, preclinical studies with follow up mechanistic experimentation are needed to confirm the causal roles of these biomarkers.

Furthermore, the availability of microbiome datasets presents obstacles for performing a more comprehensive microbiome-centric disease meta-analysis. Some diseases, such as T1D, have fewer microbiome studies, especially when compared to other conditions like IBD and CRC. Furthermore, most of the studies that we analyzed focused on a single disease, which by design excludes other disease comorbidities. At this moment, we are not aware of observational studies that investigate population-level comorbidities from a microbiome perspective. Our findings strongly suggest that broadening the range of microbiome data collection could significantly enhance the analysis of disease comorbidity. This would not only improve our understanding of known microbiome-associated diseases but could also unveil microbiome associations for disorders that have not been previously shown to have a microbiome component.

## METHODS

### Curate shotgun metagenomic datasets

The Sequence Read Archive (SRA) stands as the most extensive publicly accessible repository of sequencing data across various sequencing platforms. To identify studies and datasets exploring the human gut microbiome in the context of complex human diseases, we utilized the SRAdb package (https://github.com/seandavi/SRAdb). Using keywords such as “gut microbiome”, “human”, and “shotgun”, we identified relevant studies and datasets within the SRA repository. Datasets that have metadata available were selected and subjected to case-control matching within studies based on age and gender information. Samples were filtered to exclude individuals with obesity when BMI information was available. Unmatched samples were subsequently removed from the analysis. The retained samples then underwent consistent processing methods to generate microbial abundances. To streamline and automate the workflow, a Snakemake (79) pipeline was developed for this study, which can be accessed at (https://github.com/jindongmin/snakemake_metagenomics). The pipeline takes the SRA BioProjecct IDs as input and outputs the microbial abundance biom tables. The workflow began with downloading the sequencing data with fasterq, which was followed by quality profiling and filtering steps with fastp (80). Kraken2 (81) and bracken (82) were employed to classify the reads to the best matching location in the taxonomic tree and compute the abundance of species.

In terms of Kraken2 databases, we benchmarked Web of Life (WoL) and the Unified Human GI Genome version 2 (UHGG v2.0) databases (83). Our finding indicated that UHGG v2.0 offered a more comprehensive coverage of species at the time of our access to the databases. Consequently, we opted to utilize UHGG v2.0 in our study. Read lengths are specified based on the characteristics of the sequencing data per study.

### Build disease classifiers and filter datasets

Machine learning algorithms including GB and RF were employed for cross-validation to assess the accuracy of gut microbes in distinguishing between disease cases and control subjects. We fitted classifiers for each dataset using q2-sample-classifer (84), each microbial abundance was treated as a feature. Binary disease classifiers for CD and CRC were constructed by combining all the datasets per disease. The samples were randomly divided into training and testing sets, with an 80/20 split. The training set was utilized to construct the model and obtain optimal model parameters, while the hold-out testing dataset was used to generate predictions. The performance of GB and RF classifiers was evaluated across the datasets using the AUC. An AUC value of 0.5 indicates that the corresponding classification has the same predictive ability as random guessing. To ensure that the included datasets possessed discernible microbial signatures capable of distinguishing between cases and control subjects, we applied a threshold for AUC of 0.6 and retained only those datasets with an AUC greater than 0.6.

### SHAP interpretation of binary disease classifiers

SHAP is an explanatory approach rooted in game theory that aims to shed light on the outcomes produced by machine learning models (85). It leverages shapley values, which are a solution concept derived from cooperative game theory (86). These values provide insights into the individual contributions of players within a coalition game. In the context of SHAP, each feature (the abundance of microbes) is considered a player, and by calculating shapley values, we can ascertain their respective influences on the predictions made by disease classifiers.

### Differential abundance analysis

Microbial abundance and microbial gene abundance were analyzed using the DESeq2 package (87). The microbial species abundance data were represented as count matrices, where rows corresponded to microbial species, and columns represented samples. Healthy controls were specified as reference. Microbial species were ranked with 5% or 95% Confidence Intervals (CI) of the LFC depending on whether it decreased or increased in disease cases. And differentially abundant microbes were identified based on the ranking. Microbes with 5% CI of LFC ranked top are termed as case-associated microbes, and microbes with 95% CI of LFC ranked bottom are termed as control-associated microbes. The analysis of differential gene abundance followed a similar approach as the microbial abundance analysis. Differential gene abundances were generated with the eggNOG annotations (88). The count matrix was created with genes as rows and samples as columns. And the LFCs were calculated for further analysis.

### Disease similarity analysis

Disease similarity at the microbial species level was measured using the overlap of differentially abundant microbes. We looked at the top 100 case-associated and top 100 control-associated microbes. Concentrating on these top microbes allows us to prioritize the most relevant and significant microbial species associated with disease status. To investigate the shared microbial signatures between diseases at the species level, pairwise comparisons were conducted to determine the number of overlapping differentially abundant microbes.

Disease similarity at the microbial gene level was represented using the Pearson correlation coefficient (R) of LFCs between two diseases. A higher positive correlation coefficient value indicates a stronger similarity in the differentially abundant pattern, while a negative correlation value represents how reversed the patterns are.

### Microbial gene analysis

In order to determine if a particular gene is more commonly observed in case-associated microbes or healthy-associated microbes than by random chance, a genome wide binomial test (15) was performed between two groups of taxa. The significance level for the test was set as 0.001. Microbial genes identified that were statistically significant were subsequently mapped to KEGG pathways to elucidate their respective functional roles.

## ACKNOWLEDGEMENTS

The authors thank NYU Department of Biology and the Simons Foundation. This study used High-Performance Computing resources at the Flatiron Institute Scientific Computing Core. This work was supported by the NIAID grant 5R01AI130945.

## DATA AVAILABILITY

The following publicly available datasets were downloaded through the NCBI SRA using the following accession numbers: PRJEB47976 for Laske *et al.* (19); PRJNA451479 for Dan *et al.* (20); ERP104786 for Wang *et al.* (36); PRJNA686821 for Wan *et al.* (22); PRJNA400072 for Franzosa *et al.* (23); PRJEB15371 for a Chinese CD cohort, ERP008729 for Feng *et al.* (24); ERP005534 for Zeller *et al.* (25); PRJEB27928 for Wirbel *et al*. (26); PRJEB32762 for iMSMS Consortium. (54); PRJEB4336 for Le Chatelier *et al*. (28); ERA000116 for Qin *et al.* (*29*); PRJNA834801 for Wallen *et al.* (30); PRJNA433459 for Qian *et al.* (31); ERP111403 for Zhu *et al.* (32); PRJNA231909 for Guittar *et al.* (33); PRJNA422434 for Qin *et al*. (34); PRJEB1220 for Nielsen *et al*. (35); PRJNA400072 for Franzosa *et al.* (23); PRJEB10878 for Yu *et al.* (37). The pipeline built in this study is available at https://github.com/jindongmin/snakemake_metagenomics. The following softwares and database were used: Snakemake v 7.1.1, Bracken v2.5.2, Kraken v2.1.2, fasterq-dump v3.0.0, uhgg v2 24-Feb-2022, q2-sample-classifier v 2022.8.0, pydeseq2 v 0.2.1, shap v0.41.0.

## REFERENCE

1. Rooks MG, Garrett WS. 2016. Gut microbiota, metabolites and host immunity. Nat Rev Immunol 16:341–352.

2. Zheng D, Liwinski T, Elinav E. 2020. Interaction between microbiota and immunity in health and disease. Cell Res 30:492–506.

3. Allaband C, McDonald D, Vázquez-Baeza Y, Minich JJ, Tripathi A, Brenner DA, Loomba R, Smarr L, Sandborn WJ, Schnabl B, Dorrestein P, Zarrinpar A, Knight R. 2019. Microbiome 101: Studying, Analyzing, and Interpreting Gut Microbiome Data for Clinicians. Clin Gastroenterol Hepatol 17:218–230.

4. Rayman G, Akpan A, Cowie M, Evans R, Patel M, Posporelis S, Walsh K. 2022. Managing patients with comorbidities: future models of care. Future Healthc J 9:101–105.

5. Boersma P. 2020. Prevalence of Multiple Chronic Conditions Among US Adults, 2018. Prev Chronic Dis 17.

6. Suthram S, Dudley JT, Chiang AP, Chen R, Hastie TJ, Butte AJ. 2010. Network-based elucidation of human disease similarities reveals common functional modules enriched for pluripotent drug targets. PLoS Comput Biol 6:e1000662.

7. Clemente JC, Ursell LK, Parfrey LW, Knight R. 2012. The impact of the gut microbiota on human health: an integrative view. Cell 148:1258–1270.

8. Fan Y, Pedersen O. 2021. Gut microbiota in human metabolic health and disease. Nat Rev Microbiol 19:55–71.

9. Foster KR, Schluter J, Coyte KZ, Rakoff-Nahoum S. 2017. The evolution of the host microbiome as an ecosystem on a leash. Nature 548:43–51.

10. Maier L, Pruteanu M, Kuhn M, Zeller G, Telzerow A, Anderson EE, Brochado AR, Fernandez KC, Dose H, Mori H, Patil KR, Bork P, Typas A. 2018. Extensive impact of non-antibiotic drugs on human gut bacteria. Nature 555:623–628.

11. Balaich J, Estrella M, Wu G, Jeffrey PD, Biswas A, Zhao L, Korennykh A, Donia MS. 2021. The human microbiome encodes resistance to the antidiabetic drug acarbose. Nature 600:110–115.

12. Pianta A, Arvikar S, Strle K, Drouin EE, Wang Q, Costello CE, Steere AC. 2017. Evidence of the Immune Relevance of Prevotella copri, a Gut Microbe, in Patients With Rheumatoid Arthritis. Arthritis Rheumatol 69:964–975.

13. Pedersen HK, Gudmundsdottir V, Nielsen HB, Hyotylainen T, Nielsen T, Jensen BAH, Forslund K, Hildebrand F, Prifti E, Falony G, Le Chatelier E, Levenez F, Doré J, Mattila I, Plichta DR, Pöhö P, Hellgren LI, Arumugam M, Sunagawa S, Vieira-Silva S, Jørgensen T, Holm JB, Trošt K, MetaHIT Consortium, Kristiansen K, Brix S, Raes J, Wang J, Hansen T, Bork P, Brunak S, Oresic M, Ehrlich SD, Pedersen O. 2016. Human gut microbes impact host serum metabolome and insulin sensitivity. Nature 535:376–381.

14. Morais LH, Schreiber HL, Mazmanian SK. 2020. The gut microbiota–brain axis in behaviour and brain disorders. Nat Rev Microbiol 19:241–255.

15. Morton JT, Jin D-M, Mills RH, Shao Y, Rahman G, McDonald D, Zhu Q, Balaban M, Jiang Y, Cantrell K, Gonzalez A, Carmel J, Frankiensztajn LM, Martin-Brevet S, Berding K, Needham BD, Zurita MF, David M, Averina OV, Kovtun AS, Noto A, Mussap M, Wang M, Frank DN, Li E, Zhou W, Fanos V, Danilenko VN, Wall DP, Cárdenas P, Baldeón ME, Jacquemont S, Koren O, Elliott E, Xavier RJ, Mazmanian SK, Knight R, Gilbert JA, Donovan SM, Lawley TD, Carpenter B, Bonneau R, Taroncher-Oldenburg G. 2023. Multi-level analysis of the gut-brain axis shows autism spectrum disorder-associated molecular and microbial profiles. Nat Neurosci 26:1208–1217.

16. McLaren MR, Willis AD, Callahan BJ. 2019. Consistent and correctable bias in metagenomic sequencing experiments. Elife 8.

17. Duvallet C, Gibbons SM, Gurry T, Irizarry RA, Alm EJ. 2017. Meta-analysis of gut microbiome studies identifies disease-specific and shared responses. Nat Commun 8:1784.

18. Tierney BT, Tan Y, Kostic AD, Patel CJ. 2021. Gene-level metagenomic architectures across diseases yield high-resolution microbiome diagnostic indicators. Nat Commun 12:2907.

19. Laske C, Müller S, Preische O, Ruschil V, Munk MHJ, Honold I, Peter S, Schoppmeier U, Willmann M. 2022. Signature of Alzheimer’s Disease in Intestinal Microbiome: Results From the AlzBiom Study. Front Neurosci 16:792996.

20. Dan Z, Mao X, Liu Q, Guo M, Zhuang Y, Liu Z, Chen K, Chen J, Xu R, Tang J, Qin L, Gu B, Liu K, Su C, Zhang F, Xia Y, Hu Z, Liu X. 2020. Altered gut microbial profile is associated with abnormal metabolism activity of Autism Spectrum Disorder. Gut Microbes 11:1246– 1267.

21. Wang M, Wan J, Rong H, He F, Wang H, Zhou J, Cai C, Wang Y, Xu R, Yin Z, Zhou W. 2019. Alterations in Gut Glutamate Metabolism Associated with Changes in Gut Microbiota Composition in Children with Autism Spectrum Disorder. mSystems 4.

22. Wan Y, Zuo T, Xu Z, Zhang F, Zhan H, Chan D, Leung T-F, Yeoh YK, Chan FKL, Chan R, Ng SC. 2022. Underdevelopment of the gut microbiota and bacteria species as non-invasive markers of prediction in children with autism spectrum disorder. Gut 71:910–918.

23. Franzosa EA, Sirota-Madi A, Avila-Pacheco J, Fornelos N, Haiser HJ, Reinker S, Vatanen T, Hall AB, Mallick H, McIver LJ, Sauk JS, Wilson RG, Stevens BW, Scott JM, Pierce K, Deik AA, Bullock K, Imhann F, Porter JA, Zhernakova A, Fu J, Weersma RK, Wijmenga C, Clish CB, Vlamakis H, Huttenhower C, Xavier RJ. 2019. Gut microbiome structure and metabolic activity in inflammatory bowel disease. Nat Microbiol 4:293–305.

24. Feng Q, Liang S, Jia H, Stadlmayr A, Tang L, Lan Z, Zhang D, Xia H, Xu X, Jie Z, Su L, Li X, Li X, Li J, Xiao L, Huber-Schönauer U, Niederseer D, Xu X, Al-Aama JY, Yang H, Wang J, Kristiansen K, Arumugam M, Tilg H, Datz C, Wang J. 2015. Gut microbiome development along the colorectal adenoma-carcinoma sequence. Nat Commun 6:6528.

25. Zeller G, Tap J, Voigt AY, Sunagawa S, Kultima JR, Costea PI, Amiot A, Böhm J, Brunetti F, Habermann N, Hercog R, Koch M, Luciani A, Mende DR, Schneider MA, Schrotz-King P, Tournigand C, Tran Van Nhieu J, Yamada T, Zimmermann J, Benes V, Kloor M, Ulrich CM, von Knebel Doeberitz M, Sobhani I, Bork P. 2014. Potential of fecal microbiota for early-stage detection of colorectal cancer. Mol Syst Biol 10:766.

26. Wirbel J, Pyl PT, Kartal E, Zych K, Kashani A, Milanese A, Fleck JS, Voigt AY, Palleja A, Ponnudurai R, Sunagawa S, Coelho LP, Schrotz-King P, Vogtmann E, Habermann N, Niméus E, Thomas AM, Manghi P, Gandini S, Serrano D, Mizutani S, Shiroma H, Shiba S, Shibata T, Yachida S, Yamada T, Waldron L, Naccarati A, Segata N, Sinha R, Ulrich CM, Brenner H, Arumugam M, Bork P, Zeller G. 2019. Meta-analysis of fecal metagenomes reveals global microbial signatures that are specific for colorectal cancer. Nat Med 25:679– 689.

27. The iMSMS Consortium. 2020. Household paired design reduces variance and increases power in multi-city gut microbiome study in multiple sclerosis. Mult Scler 1352458520924594.

28. Le Chatelier E, Nielsen T, Qin J, Prifti E, Hildebrand F, Falony G, Almeida M, Arumugam M, Batto J-M, Kennedy S, Leonard P, Li J, Burgdorf K, Grarup N, Jørgensen T, Brandslund I, Nielsen HB, Juncker AS, Bertalan M, Levenez F, Pons N, Rasmussen S, Sunagawa S, Tap J, Tims S, Zoetendal EG, Brunak S, Clément K, Doré J, Kleerebezem M, Kristiansen K, Renault P, Sicheritz-Ponten T, de Vos WM, Zucker J-D, Raes J, Hansen T, MetaHIT consortium, Bork P, Wang J, Ehrlich SD, Pedersen O. 2013. Richness of human gut microbiome correlates with metabolic markers. Nature 500:541–546.

29. Qin J, Li R, Raes J, Arumugam M, Burgdorf KS, Manichanh C, Nielsen T, Pons N, Levenez F, Yamada T, Mende DR, Li J, Xu J, Li S, Li D, Cao J, Wang B, Liang H, Zheng H, Xie Y, Tap J, Lepage P, Bertalan M, Batto J-M, Hansen T, Le Paslier D, Linneberg A, Nielsen HB, Pelletier E, Renault P, Sicheritz-Ponten T, Turner K, Zhu H, Yu C, Li S, Jian M, Zhou Y, Li Y, Zhang X, Li S, Qin N, Yang H, Wang J, Brunak S, Doré J, Guarner F, Kristiansen K, Pedersen O, Parkhill J, Weissenbach J, MetaHIT Consortium, Bork P, Ehrlich SD, Wang J. 2010. A human gut microbial gene catalogue established by metagenomic sequencing. Nature 464:59–65.

30. Wallen ZD, Demirkan A, Twa G, Cohen G, Dean MN, Standaert DG, Sampson TR, Payami H. 2022. Metagenomics of Parkinson’s disease implicates the gut microbiome in multiple disease mechanisms. Nat Commun 13:6958.

31. Qian Y, Yang X, Xu S, Huang P, Li B, Du J, He Y, Su B, Xu L-M, Wang L, Huang R, Chen S, Xiao Q. 2020. Gut metagenomics-derived genes as potential biomarkers of Parkinson’s disease. Brain 143:2474–2489.

32. Zhu F, Ju Y, Wang W, Wang Q, Guo R, Ma Q, Sun Q, Fan Y, Xie Y, Yang Z, Jie Z, Zhao B, Xiao L, Yang L, Zhang T, Feng J, Guo L, He X, Chen Y, Chen C, Gao C, Xu X, Yang H, Wang J, Dang Y, Madsen L, Brix S, Kristiansen K, Jia H, Ma X. 2020. Metagenome-wide association of gut microbiome features for schizophrenia. Nat Commun 11:1612.

33. Guittar J, Shade A, Litchman E. 2019. Trait-based community assembly and succession of the infant gut microbiome. Nature Communications 10.1038/s41467-019-08377-w.

34. Qin J, Li Y, Cai Z, Li S, Zhu J, Zhang F, Liang S, Zhang W, Guan Y, Shen D, Peng Y, Zhang D, Jie Z, Wu W, Qin Y, Xue W, Li J, Han L, Lu D, Wu P, Dai Y, Sun X, Li Z, Tang A, Zhong S, Li X, Chen W, Xu R, Wang M, Feng Q, Gong M, Yu J, Zhang Y, Zhang M, Hansen T, Sanchez G, Raes J, Falony G, Okuda S, Almeida M, LeChatelier E, Renault P, Pons N, Batto J-M, Zhang Z, Chen H, Yang R, Zheng W, Li S, Yang H, Wang J, Ehrlich SD, Nielsen R, Pedersen O, Kristiansen K, Wang J. 2012. A metagenome-wide association study of gut microbiota in type 2 diabetes. Nature 490:55–60.

35. Nielsen HB, Almeida M, Juncker AS, Rasmussen S, Li J, Sunagawa S, Plichta DR, Gautier L, Pedersen AG, Le Chatelier E, Pelletier E, Bonde I, Nielsen T, Manichanh C, Arumugam M, Batto J-M, Quintanilha Dos Santos MB, Blom N, Borruel N, Burgdorf KS, Boumezbeur F, Casellas F, Doré J, Dworzynski P, Guarner F, Hansen T, Hildebrand F, Kaas RS, Kennedy S, Kristiansen K, Kultima JR, Léonard P, Levenez F, Lund O, Moumen B, Le Paslier D, Pons N, Pedersen O, Prifti E, Qin J, Raes J, Sørensen S, Tap J, Tims S, Ussery DW, Yamada T, MetaHIT Consortium, Renault P, Sicheritz-Ponten T, Bork P, Wang J, Brunak S, Ehrlich SD, MetaHIT Consortium. 2014. Identification and assembly of genomes and genetic elements in complex metagenomic samples without using reference genomes. Nat Biotechnol 32:822–828.

36. Wang M, Doenyas C, Wan J, Zeng S, Cai C, Zhou J, Liu Y, Yin Z, Zhou W. 2021. Virulence factor-related gut microbiota genes and immunoglobulin A levels as novel markers for machine learning-based classification of autism spectrum disorder. Comput Struct Biotechnol J 19:545–554.

37. Yu J, Feng Q, Wong SH, Zhang D, Liang QY, Qin Y, Tang L, Zhao H, Stenvang J, Li Y, Wang X, Xu X, Chen N, Wu WKK, Al-Aama J, Nielsen HJ, Kiilerich P, Jensen BAH, Yau TO, Lan Z, Jia H, Li J, Xiao L, Lam TYT, Ng SC, Cheng AS-L, Wong VW-S, Chan FKL, Xu X, Yang H, Madsen L, Datz C, Tilg H, Wang J, Brünner N, Kristiansen K, Arumugam M, Sung JJ-Y, Wang J. 2017. Metagenomic analysis of faecal microbiome as a tool towards targeted non-invasive biomarkers for colorectal cancer. Gut 66:70–78.

38. Ullman TA, Itzkowitz SH. 2011. Intestinal inflammation and cancer. Gastroenterology 140:1807–1816.

39. Quévrain E, Maubert MA, Michon C, Chain F, Marquant R, Tailhades J, Miquel S, Carlier L, Bermúdez-Humarán LG, Pigneur B, Lequin O, Kharrat P, Thomas G, Rainteau D, Aubry C, Breyner N, Afonso C, Lavielle S, Grill J-P, Chassaing G, Chatel JM, Trugnan G, Xavier R, Langella P, Sokol H, Seksik P. 2016. Identification of an anti-inflammatory protein from Faecalibacterium prausnitzii, a commensal bacterium deficient in Crohn’s disease. Gut 65:415–425.

40. Abed J, Maalouf N, Manson AL, Earl AM, Parhi L, Emgård JEM, Klutstein M, Tayeb S, Almogy G, Atlan KA, Chaushu S, Israeli E, Mandelboim O, Garrett WS, Bachrach G. 2020. Colon Cancer-Associated May Originate From the Oral Cavity and Reach Colon Tumors via the Circulatory System. Front Cell Infect Microbiol 10:400.

41. 2013. Fusobacterium nucleatum Potentiates Intestinal Tumorigenesis and Modulates the Tumor-Immune Microenvironment. Cell Host Microbe 14:207–215.

42. Segal LN, Clemente JC, Tsay J-CJ, Koralov SB, Keller BC, Wu BG, Li Y, Shen N, Ghedin E, Morris A, Diaz P, Huang L, Wikoff WR, Ubeda C, Artacho A, Rom WN, Sterman DH, Collman RG, Blaser MJ, Weiden MD. 2016. Enrichment of the lung microbiome with oral taxa is associated with lung inflammation of a Th17 phenotype. Nat Microbiol 1:16031.

43. Tsay J-CJ, Wu BG, Sulaiman I, Gershner K, Schluger R, Li Y, Yie T-A, Meyn P, Olsen E, Perez L, Franca B, Carpenito J, Iizumi T, El-Ashmawy M, Badri M, Morton JT, Shen N, He L, Michaud G, Rafeq S, Bessich JL, Smith RL, Sauthoff H, Felner K, Pillai R, Zavitsanou A- M, Koralov SB, Mezzano V, Loomis CA, Moreira AL, Moore W, Tsirigos A, Heguy A, Rom WN, Sterman DH, Pass HI, Clemente JC, Li H, Bonneau R, Wong K-K, Papagiannakopoulos T, Segal LN. 2021. Lower Airway Dysbiosis Affects Lung Cancer Progression. Cancer Discov 11:293–307.

44. Kitamoto S, Nagao-Kitamoto H, Jiao Y, Gillilland MG 3rd, Hayashi A, Imai J, Sugihara K, Miyoshi M, Brazil JC, Kuffa P, Hill BD, Rizvi SM, Wen F, Bishu S, Inohara N, Eaton KA, Nusrat A, Lei YL, Giannobile WV, Kamada N. 2020. The Intermucosal Connection between the Mouth and Gut in Commensal Pathobiont-Driven Colitis. Cell 182:447–462.e14.

45. Lutgens MWMD, van Oijen MGH, van der Heijden GJMG, Vleggaar FP, Siersema PD, Oldenburg B. 2013. Declining risk of colorectal cancer in inflammatory bowel disease: an updated meta-analysis of population-based cohort studies. Inflamm Bowel Dis 19:789–799.

46. Jang H-M, Kim J-K, Joo M-K, Shin Y-J, Lee K-E, Lee CK, Kim H-J, Kim D-H. 2022. Enterococcus faecium and Pediococcus acidilactici deteriorate Enterobacteriaceae-induced depression and colitis in mice. Sci Rep 12:9389.

47. Cao Y, Oh J, Xue M, Huh WJ, Wang J, Gonzalez-Hernandez JA, Rice TA, Martin AL, Song D, Crawford JM, Herzon SB, Palm NW. 2022. Commensal microbiota from patients with inflammatory bowel disease produce genotoxic metabolites. Science 378:eabm3233.

48. Yang R, Shan S, Shi J, Li H, An N, Li S, Cui K, Guo H, Li Z. 2023. , a Potent Probiotic, Alleviates Colitis via Acetate-Mediated IgA Response and Microbiota Restoration. J Agric Food Chem 10.1021/acs.jafc.2c06697.

49. Ma S, Shungin D, Mallick H, Schirmer M, Nguyen LH, Kolde R, Franzosa E, Vlamakis H, Xavier R, Huttenhower C. 2022. Population structure discovery in meta-analyzed microbial communities and inflammatory bowel disease using MMUPHin. Genome Biol 23:208.

50. Ruuskanen MO, Erawijantari PP, Havulinna AS, Liu Y, Méric G, Tuomilehto J, Inouye M, Jousilahti P, Salomaa V, Jain M, Knight R, Lahti L, Niiranen TJ. 2022. Gut Microbiome Composition Is Predictive of Incident Type 2 Diabetes in a Population Cohort of 5,572 Finnish Adults. Diabetes Care 45:811–818.

51. McGuinness AJ, Davis JA, Dawson SL, Loughman A, Collier F, O’Hely M, Simpson CA, Green J, Marx W, Hair C, Guest G, Mohebbi M, Berk M, Stupart D, Watters D, Jacka FN. 2022. A systematic review of gut microbiota composition in observational studies of major depressive disorder, bipolar disorder and schizophrenia. Mol Psychiatry 27:1920–1935.

52. Zheng J, Hoffman KL, Chen J-S, Shivappa N, Sood A, Browman GJ, Dirba DD, Hanash S, Wei P, Hebert JR, Petrosino JF, Schembre SM, Daniel CR. 2020. Dietary inflammatory potential in relation to the gut microbiome: results from a cross-sectional study. Br J Nutr 124:931–942.

53. Xue C, Li G, Gu X, Su Y, Zheng Q, Yuan X, Bao Z, Lu J, Li L. 2023. Health and Disease:, the Shining Star of the Gut Flora. Research 6:0107.

54. iMSMS Consortium. Electronic address, iMSMS Consortium. 2022. Gut microbiome of multiple sclerosis patients and paired household healthy controls reveal associations with disease risk and course. Cell 185:3467–3486.e16.

55. Lopez J, Grinspan A. 2016. Fecal Microbiota Transplantation for Inflammatory Bowel Disease. Gastroenterol Hepatol 12:374–379.

56. Hellmig S, Ott S, Musfeldt M, Kosmahl M, Rosenstiel P, Stüber E, Hampe J, Fölsch UR, Schreiber S. 2005. Life-threatening chronic enteritis due to colonization of the small bowel with Stenotrophomonas maltophilia. Gastroenterology 129:706–712.

57. Rashid T, Ebringer A, Wilson C. 2013. The role of Klebsiella in Crohn’s disease with a potential for the use of antimicrobial measures. Int J Rheumatol 2013:610393.

58. Liu H, Hong XL, Sun TT, Huang XW, Wang JL, Xiong H. 2020. Fusobacterium nucleatum exacerbates colitis by damaging epithelial barriers and inducing aberrant inflammation. J Dig Dis 21:385–398.

59. Das TK, Ganesh BP. 2023. Interlink between the gut microbiota and inflammation in the context of oxidative stress in Alzheimer’s disease progression. Gut Microbes 15:2206504.

60. Xu L, Liu R, Qin Y, Wang T. 2023. Brain metabolism in Alzheimer’s disease: biological mechanisms of exercise. Transl Neurodegener 12:33.

61. van Dyck CH, Swanson CJ, Aisen P, Bateman RJ, Chen C, Gee M, Kanekiyo M, Li D, Reyderman L, Cohen S, Froelich L, Katayama S, Sabbagh M, Vellas B, Watson D, Dhadda S, Irizarry M, Kramer LD, Iwatsubo T. 2023. Lecanemab in Early Alzheimer’s Disease. N Engl J Med 388:9–21.

62. Chande N, Townsend CM, Parker CE, MacDonald JK. 2016. Azathioprine or 6-mercaptopurine for induction of remission in Crohn’s disease. Cochrane Database Syst Rev 10:CD000545.

63. Gevers D, Kugathasan S, Denson LA, Vázquez-Baeza Y, Van Treuren W, Ren B, Schwager E, Knights D, Song SJ, Yassour M, Morgan XC, Kostic AD, Luo C, González A, McDonald D, Haberman Y, Walters T, Baker S, Rosh J, Stephens M, Heyman M, Markowitz J, Baldassano R, Griffiths A, Sylvester F, Mack D, Kim S, Crandall W, Hyams J, Huttenhower C, Knight R, Xavier RJ. 2014. The treatment-naive microbiome in new-onset Crohn’s disease. Cell Host Microbe 15:382–392.

64. Human Microbiome Project Consortium. 2012. Structure, function and diversity of the healthy human microbiome. Nature 486:207–214.

65. Lee M-K, Lee S-Y, Sohn S-Y, Ahn J, Han K, Lee J-H. 2023. Type 2 Diabetes and Its Association With Psychiatric Disorders in Young Adults in South Korea. JAMA Netw Open 6:e2319132.

66. Rojas LBA, Gomes MB. 2013. Metformin: an old but still the best treatment for type 2 diabetes. Diabetol Metab Syndr 5:6.

67. Wu H, Esteve E, Tremaroli V, Khan MT, Caesar R, Mannerås-Holm L, Ståhlman M, Olsson LM, Serino M, Planas-Fèlix M, Xifra G, Mercader JM, Torrents D, Burcelin R, Ricart W, Perkins R, Fernàndez-Real JM, Bäckhed F. 2017. Metformin alters the gut microbiome of individuals with treatment-naive type 2 diabetes, contributing to the therapeutic effects of the drug. Nat Med 23:850–858.

68. Zhang Y-M, Zong H-C, Qi Y-B, Chang L-L, Gao Y-N, Zhou T, Yin T, Liu M, Pan K-J, Chen W-G, Guo H-R, Guo F, Peng Y-M, Wang M, Feng L-Y, Zang Y, Li Y, Li J. 2023. Anxiolytic effect of antidiabetic metformin is mediated by AMPK activation in mPFC inhibitory neurons. Mol Psychiatry 10.1038/s41380-023-02283-w.

69. Kessing LV, Rytgaard HC, Ekstrøm CT, Knop FK, Berk M, Gerds TA. 2020. Antidiabetes Agents and Incident Depression: A Nationwide Population-Based Study. Diabetes Care 43:3050–3060.

70. Patil SP, Jain PD, Ghumatkar PJ, Tambe R, Sathaye S. 2014. Neuroprotective effect of metformin in MPTP-induced Parkinson’s disease in mice. Neuroscience 277:747–754.

71. Paudel YN, Angelopoulou E, Piperi C, Shaikh MF, Othman I. 2020. Emerging neuroprotective effect of metformin in Parkinson’s disease: A molecular crosstalk. Pharmacol Res 152:104593.

72. Grabrucker S, Marizzoni M, Silajdžić E, Lopizzo N, Mombelli E, Nicolas S, Dohm-Hansen S, Scassellati C, Moretti DV, Rosa M, Hoffmann K, Cryan JF, O’Leary OF, English JA, Lavelle A, O’Neill C, Thuret S, Cattaneo A, Nolan YM. 2023. Microbiota from Alzheimer’s patients induce deficits in cognition and hippocampal neurogenesis. Brain 10.1093/brain/awad303.

73. Kedia S, Virmani S, K Vuyyuru S, Kumar P, Kante B, Sahu P, Kaushal K, Farooqui M, Singh M, Verma M, Bajaj A, Markandey M, Sachdeva K, Das P, Makharia GK, Ahuja V. 2022. Faecal microbiota transplantation with anti-inflammatory diet (FMT-AID) followed by anti-inflammatory diet alone is effective in inducing and maintaining remission over 1 year in mild to moderate ulcerative colitis: a randomised controlled trial. Gut 71:2401–2413.

74. Pigneur B, Sokol H. 2016. Fecal microbiota transplantation in inflammatory bowel disease: the quest for the holy grail. Mucosal Immunol 9:1360–1365.

75. Morton JT, Marotz C, Washburne A, Silverman J, Zaramela LS, Edlund A, Zengler K, Knight R. 2019. Establishing microbial composition measurement standards with reference frames. Nat Commun 10:2719.

76. Nichols RG, Davenport ER. 2021. The relationship between the gut microbiome and host gene expression: a review. Hum Genet 140:747–760.

77. Priya S, Burns MB, Ward T, Mars RAT, Adamowicz B, Lock EF, Kashyap PC, Knights D, Blekhman R. 2022. Identification of shared and disease-specific host gene-microbiome associations across human diseases using multi-omic integration. Nat Microbiol 7:780–795.

78. Quinn RA, Melnik AV, Vrbanac A, Fu T, Patras KA, Christy MP, Bodai Z, Belda-Ferre P, Tripathi A, Chung LK, Downes M, Welch RD, Quinn M, Humphrey G, Panitchpakdi M, Weldon KC, Aksenov A, da Silva R, Avila-Pacheco J, Clish C, Bae S, Mallick H, Franzosa EA, Lloyd-Price J, Bussell R, Thron T, Nelson AT, Wang M, Leszczynski E, Vargas F, Gauglitz JM, Meehan MJ, Gentry E, Arthur TD, Komor AC, Poulsen O, Boland BS, Chang JT, Sandborn WJ, Lim M, Garg N, Lumeng JC, Xavier RJ, Kazmierczak BI, Jain R, Egan M, Rhee KE, Ferguson D, Raffatellu M, Vlamakis H, Haddad GG, Siegel D, Huttenhower C, Mazmanian SK, Evans RM, Nizet V, Knight R, Dorrestein PC. 2020. Global chemical effects of the microbiome include new bile-acid conjugations. Nature 579:123–129.

79. Köster J, Rahmann S. 2018. Snakemake-a scalable bioinformatics workflow engine. Bioinformatics 34:3600.

80. Chen S, Zhou Y, Chen Y, Gu J. 2018. fastp: an ultra-fast all-in-one FASTQ preprocessor. Bioinformatics 34:i884–i890.

81. Wood DE, Lu J, Langmead B. 2019. Improved metagenomic analysis with Kraken 2. Genome Biol 20:257.

82. Lu J, Breitwieser FP, Thielen P, Salzberg SL. 2017. Bracken: estimating species abundance in metagenomics data. PeerJ Comput Sci 3:e104.

83. Almeida A, Nayfach S, Boland M, Strozzi F, Beracochea M, Shi ZJ, Pollard KS, Sakharova E, Parks DH, Hugenholtz P, Segata N, Kyrpides NC, Finn RD. 2021. A unified catalog of 204,938 reference genomes from the human gut microbiome. Nat Biotechnol 39:105–114.

84. Bokulich NA, Dillon MR, Bolyen E, Kaehler BD, Huttley GA, Caporaso JG. 2018. q2-sample-classifier: machine-learning tools for microbiome classification and regression. J Open Res Softw 3.

85. Lundberg SM, Erion G, Chen H, DeGrave A, Prutkin JM, Nair B, Katz R, Himmelfarb J, Bansal N, Lee S-I. 2020. From Local Explanations to Global Understanding with Explainable AI for Trees. Nat Mach Intell 2:56–67.

86. Shapley LS. 1952. A Value for N-person Games.

87. Love MI, Huber W, Anders S. 2014. Moderated estimation of fold change and dispersion for RNA-seq data with DESeq2. Genome Biol 15:550.

88. Cantalapiedra CP, Hernández-Plaza A, Letunic I, Bork P, Huerta-Cepas J. 2021. eggNOG-mapper v2: Functional Annotation, Orthology Assignments, and Domain Prediction at the Metagenomic Scale. Mol Biol Evol 38:5825–5829.

